# Theta-Band Temporal Interference Stimulation Targeting the dACC Modulates Neural Gain of Prediction Error Encoding

**DOI:** 10.64898/2025.12.20.695275

**Authors:** Yatong Wen, Mo Wang, Xinwen Dong, Mateusz Gola, Quanying Liu, Yiheng Tu, Yonghui Li

**Author notes:** Correspondence: Yiheng Tu, State Key Laboratory of Cognitive Science and Mental Health, Institute of Psychology, Chinese Academy of Sciences, No.16 Lincui Road, Chaoyang District, Beijing 100101, China. and Yonghui Li, State Key Laboratory of Cognitive Science and Mental Health, Institute of Psychology, Chinese Academy of Sciences, No.16 Lincui Road, Chaoyang District, Beijing 100101, China.

## Abstract

**Background:** Prediction error (PE) signals encode discrepancies between expected and actual outcomes and are linked to medial frontal theta dynamics in the dorsal anterior cingulate cortex (dACC). However, causal evidence that dACC theta mechanisms contribute to PE gain in humans remains limited. Transcranial temporal interference stimulation (tTIS) enables non-invasive modulation of deep cortical targets and provides a strategy to probe dACC theta circuitry.

**Objective:** To determine whether theta-band tTIS targeting the dACC modulates the neural gain of prediction error encoding during reward processing.

**Methods:** Thirty-four participants with elevated food addiction-related symptoms were randomized in a single-blind, sham-controlled pre-post design to active theta-band tTIS or sham stimulation. Under satiety, participants completed a food incentive delay task while EEG was recorded before and after stimulation. Feedback-related negativity (FRN) was analyzed at mean and single-trial levels using mixed-effects models with behavioral and Rescorla-Wagner model-derived prediction errors. Stimulus-preceding negativity (SPN) indexed anticipatory processing.

**Results:** Theta-band tTIS selectively modulated feedback-stage prediction error encoding. It altered the trial-level FRN–prediction error slope, with pre-to-post changes differing between groups for both behavioral (*β* = 1.74, *p* = .049) and model-derived prediction errors (*β* = 1.77, *p* = .048). At the mean ERP level, FRN became more negative after active stimulation (*t*_16_ = 2.75, *p* = .014), whereas SPN was unaffected.

**Conclusion:** Theta-band tTIS targeting the dACC modulates neural sensitivity to prediction errors during reward feedback, providing causal evidence that medial frontal theta dynamics regulate the neural gain of prediction error encoding.

## Introduction

Prediction error signals guide adaptive learning by encoding discrepancies between expected and actual outcomes [1–3]. Converging electrophysiological and neuroimaging evidence indicates that medial frontal theta-band dynamics have been implicated in this process [4], with the dorsal anterior cingulate cortex (dACC) serving as a key generator of feedback-related prediction error activity [5]. Neuronal recordings demonstrate that dACC ensembles dynamically update firing patterns in response to outcome deviations [6], whereas synchronized theta oscillations are thought to coordinate performance monitoring across prefrontal networks [4] and support adaptive behavioral adjustments [7]. Together, these converging neuronal and oscillatory findings implicate dACC theta dynamics as a candidate mechanism regulating the neural gain of prediction error encoding in humans. However, causal tests of these dACC-centered, frequency-specific oscillatory mechanisms in humans remain limited.

Conventional non-invasive stimulation approaches provide limited access to deep medial frontal cortex such as the dACC. Transcranial alternating current stimulation (tACS) demonstrates frequency-dependent modulation of cortical rhythms [8–10]. Phase- and state-dependent stimulation can also modulate oscillatory dynamics and influence neural gain in a behaviorally relevant manner [11,12]. However, these approaches predominantly influence superficial cortical regions and are poorly suited for selectively engaging deeper medial frontal structures. Invasive recordings obtained during deep brain stimulation (DBS) procedures provide rare human evidence linking dACC activity to prediction error processing [13]. However, DBS is confined to clinical contexts and does not permit controlled frequency-specific interrogation of oscillatory mechanisms. A non-invasive method capable of selectively modulating deep cortical oscillations at a defined frequency would therefore provide critical mechanistic insight into whether medial frontal theta dynamics regulate prediction error gain. Transcranial temporal interference stimulation (tTIS) enables non-invasive modulation of deep brain regions by generating a low-frequency envelope through the interaction of two high-frequency currents [14]. Biophysical modeling suggests that temporal interference can concentrate envelope modulation in deep structures while minimizing stimulation of overlying cortex [15]. Emerging human evidence further supports its capacity to modulate deep oscillatory circuits [16,17]. This approach enables causal testing of deep, frequency-specific modulation of dACC activity during prediction error encoding.

Here, we tested whether theta-band tTIS targeting the dACC modulates the neural gain of prediction error encoding during reward processing. The experiment was conducted in individuals exhibiting elevated reward expectancy bias, operationalized as elevated food addiction-related symptoms assessed using validated questionnaires [18,19]. Prior work indicates that this expectancy bias persists even under satiety and is associated with reduced sensitivity to outcome discrepancies [18]. Such persistent expectancy bias provides a model in which deviations between expected and actual outcomes may be informative for probing prediction error dynamics. In this context, unfavorable (negative) prediction errors provide a sensitive index of updating; analyses therefore focused on negative prediction error trials. We hypothesized that theta-band deep modulation would alter the coupling between electrophysiological indices of dACC activity and trial-by-trial prediction errors.

## Materials and methods

### Participants

Thirty-four participants with elevated food addiction-related symptoms were recruited (15 males, 19 females). Symptoms were assessed using the Yale Food Addiction Scale (YFAS), with inclusion criteria of a YFAS symptom score ≥ 3 and BMI ≥ 25, consistent with prior studies [20–22]. The mean age of the sample was 25.32 ± 4.64 years (range: 19–39 years), and the mean body mass index (BMI) was 27.75 ± 5.05 kg/m². Exclusion criteria included a history of neurological or psychiatric disorders, current use of psychoactive medication, epilepsy or seizure history, metal implants in the head, dermatological conditions affecting the scalp, pregnancy, or contraindications to non-invasive brain stimulation.

All participants had normal or corrected-to-normal vision and provided written informed consent in accordance with the Declaration of Helsinki. The study protocol was approved by the Ethics Committee of the Institute of Psychology, Chinese Academy of Sciences on October 31, 2023 (No. H23113).

Participants were randomly assigned to the active tTIS group (n = 17) or the sham stimulation group (n = 17). Participants consumed lunch before testing and refrained from further caloric intake. Hunger levels were assessed immediately before the experiment using a 10-point visual scale, and all participants reported hunger ratings below 5. Participants received a base payment of ¥200–¥230, with an additional performance-based bonus of up to ¥15.

### Study design and procedure

The study employed a mixed 2 × 2 design with Group (active tTIS vs. sham stimulation) as a between-subject factor and Phase (pre-vs. post-stimulation) as a within-subject factor. The experiment comprised pretest, stimulation, and posttest phases. EEG was recorded during the 150-trial pretest and posttest, but not during stimulation. Before the main experiment, participants completed a practice session to familiarize themselves with the task and were required to achieve at least 80% accuracy before proceeding. All trials involved food rewards represented by standardized food images matched on arousal, pleasantness, and familiarity (Supplementary Section 1; Table S1). The overall experimental timeline is illustrated in Figure 1, and the task structure is shown in Figure 2.

**Figure 1.**
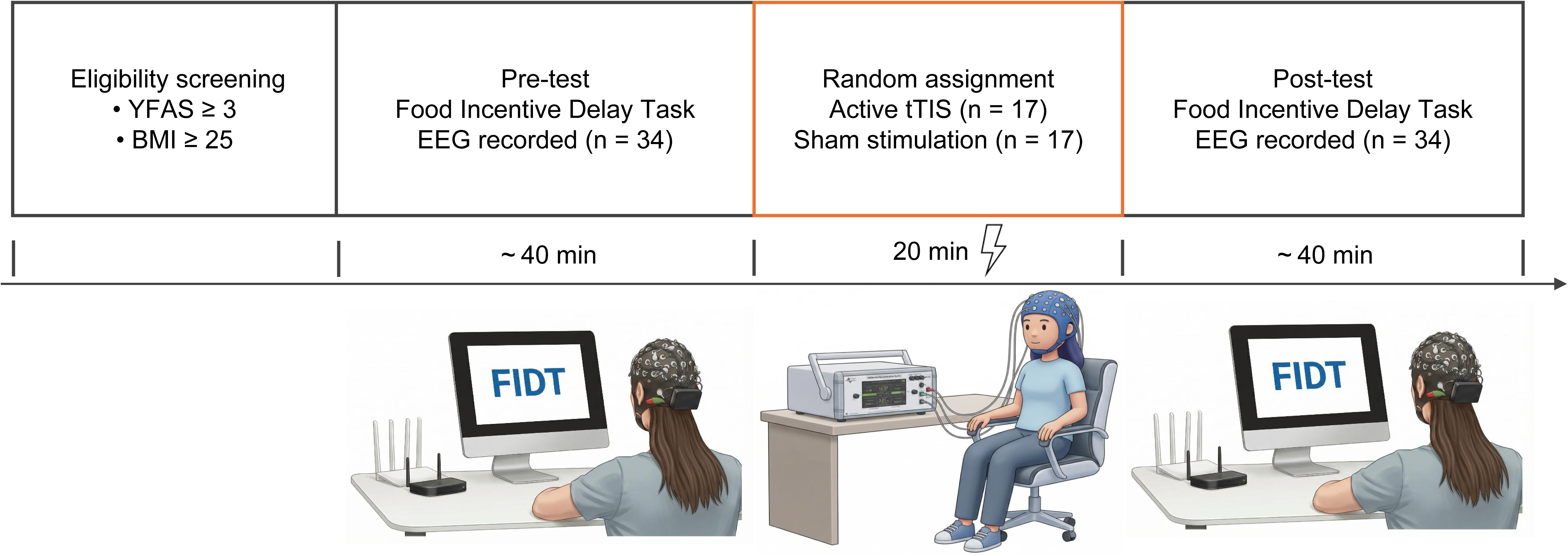
Experimental design and procedure. Eligible participants proceeded to a pre-test session of a food incentive delay task with concurrent EEG recording. Participants were then randomly assigned to receive either active theta-band transcranial temporal interference stimulation (tTIS; n = 17) or sham stimulation (n = 17) for 20 minutes. Following stimulation, all participants completed a post-test session identical to the pre-test. Both pre-test and post-test sessions lasted approximately 40 minutes.

**Figure 2.**
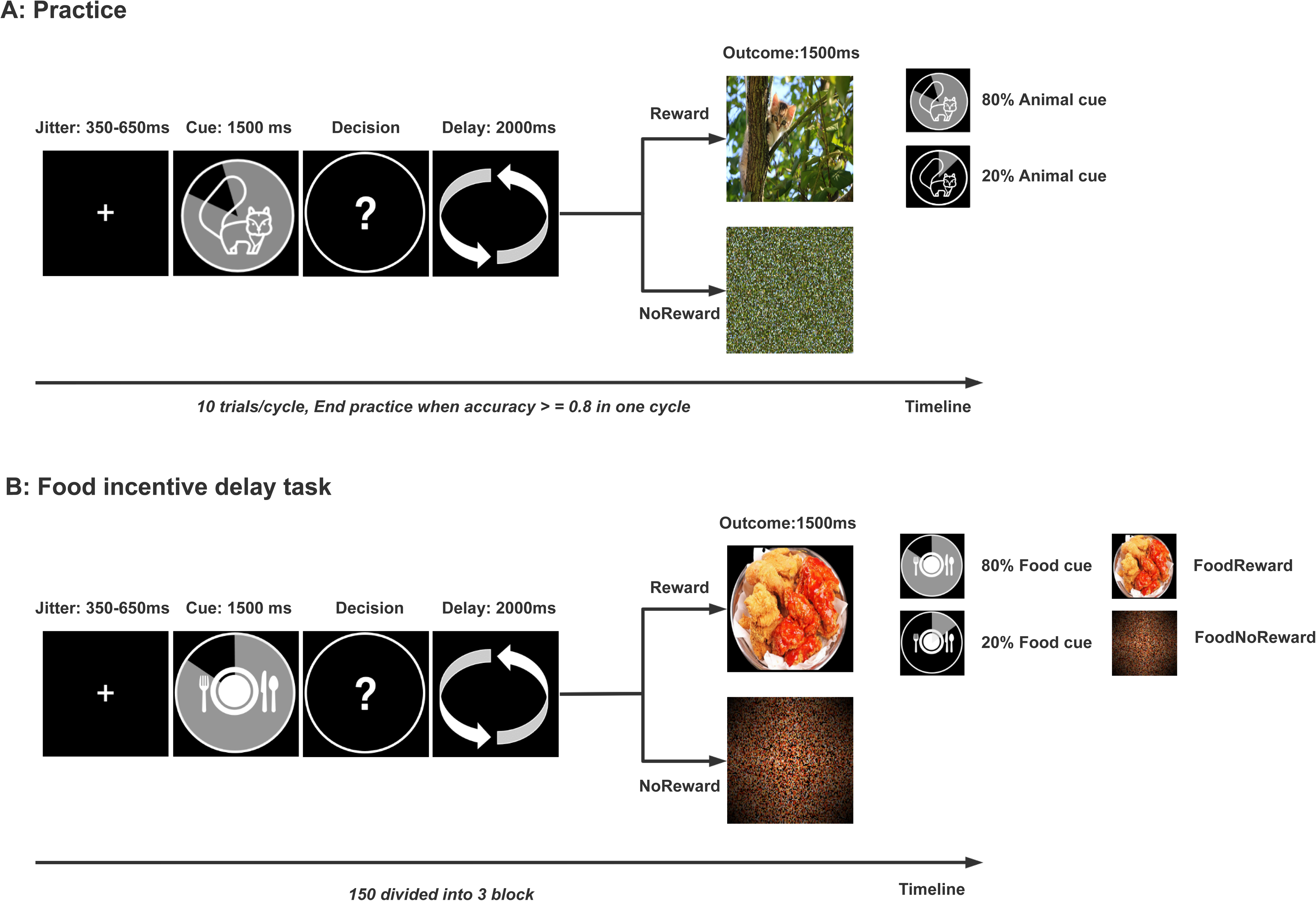
Experimental task design scheme. (A) Practice session. Participants completed a brief practice version of the incentive delay task using natural reward stimuli. Practice ended when accuracy reached ≥ 0.80 within one cycle. (B) Food incentive delay task. Each trial began with a fixation cross followed by a food cue (1500 ms) indicating reward probability (80% high vs. 20% low). Participants then reported reward expectation via key press (self-paced), followed by a 2000-ms delay and outcome presentation (1500 ms; food image vs. scrambled image). Inter-trial intervals were jittered between 350 and 650 ms. Each pre- and post-stimulation phase consisted of 150 trials, divided into three blocks of 50 trials with short breaks between blocks.

### Incentive Delay Task

Reward processing was assessed using a modified Incentive Delay Task (IDT) adapted from Knutson et al. [23], focusing on food reward outcomes. Each trial began with a cue indicating the probability of receiving a food reward (high: 80%; low: 20%), followed by a response phase in which participants indicated whether they expected to receive a reward (yes/no) using a binary button press (“F” or “J”). After a 2000-ms delay, feedback was presented for 1500 ms. Reward feedback consisted of the presentation of a food image (reward) or a scrambled image (no reward). Inter-trial intervals varied between 350 and 650 ms. Behavioral outcomes included reaction times and reward prediction rates under high- and low-probability conditions. These trial-wise expectation–outcome combinations were used to derive behavioral prediction errors coded as −1 (worse-than-expected), 0 (expected), and +1 (better-than-expected).

### Electroencephalogram (EEG) recording and processing

Continuous EEG was recorded using a 64-channel system (Neuracle NeuSen.W 64; Neuracle, Changzhou, China), referenced to CPz with AFz as ground. Signals were sampled at 1000 Hz, electrode impedances were kept below 10 kΩ, and data were rereferenced offline to the average of the bilateral mastoids (M1/M2). Analyses focused on two ERP components: feedback-related negativity (FRN), indexing feedback evaluation, and stimulus-preceding negativity (SPN), indexing anticipatory processing.

EEG preprocessing was conducted in EEGLAB/ERPLAB using custom MATLAB scripts. Data were band-pass filtered using zero-phase finite impulse response filters (FRN: 0.1–30 Hz; SPN: low-pass filter at 30 Hz), epoched relative to feedback onset, and cleaned using automated rejection, manual inspection, and independent component analysis (ICA; runica algorithm) to remove ocular and muscle artifacts. Trials with residual artifacts exceeding ±100 μV were automatically rejected. SPN epochs were extracted from −2000 to 500 ms (baseline: −2000 to −1800 ms), and FRN epochs from −200 to 1000 ms (baseline: −200 to 0 ms). Single-trial amplitudes were extracted for mixed-effects analyses, and grand averages were computed for visualization.

### Transcranial Temporal Interference Stimulation

tTIS was delivered using an interferential neuromodulation system (Soterix Medical Inc.) targeting the dACC. Ag/AgCl electrodes (1.2 cm diameter; contact area = 1.13 cm²) were placed according to an optimized unilateral montage [24] (Supplementary Section 2). Target coordinates (MNI: −5, 34, 30) were derived from a prior neuromodulation study [25] and verified using FSL and MNI–Talairach conversion tools (see URL in Supplementary Section 2). Electrodes were attached using conductive gel and secured to ensure stable contact.

Four electrodes were used (AF3 positive, Fz negative, CP3 positive, F7 negative). Finite-element simulations estimated the temporal interference envelope distribution and confirmed field intensity in the dACC target (Figure 3). Stimulation used two high-frequency currents (2000 and 2005 Hz) producing a 5-Hz theta-band envelope. Current intensity was 1.6 mA (peak-to-baseline) per electrode for 20 minutes. Active stimulation included gradual ramp-in and ramp-out periods, whereas sham stimulation consisted of a 30-s ramp-up and ramp-down without sustained stimulation.

**Figure 3.**
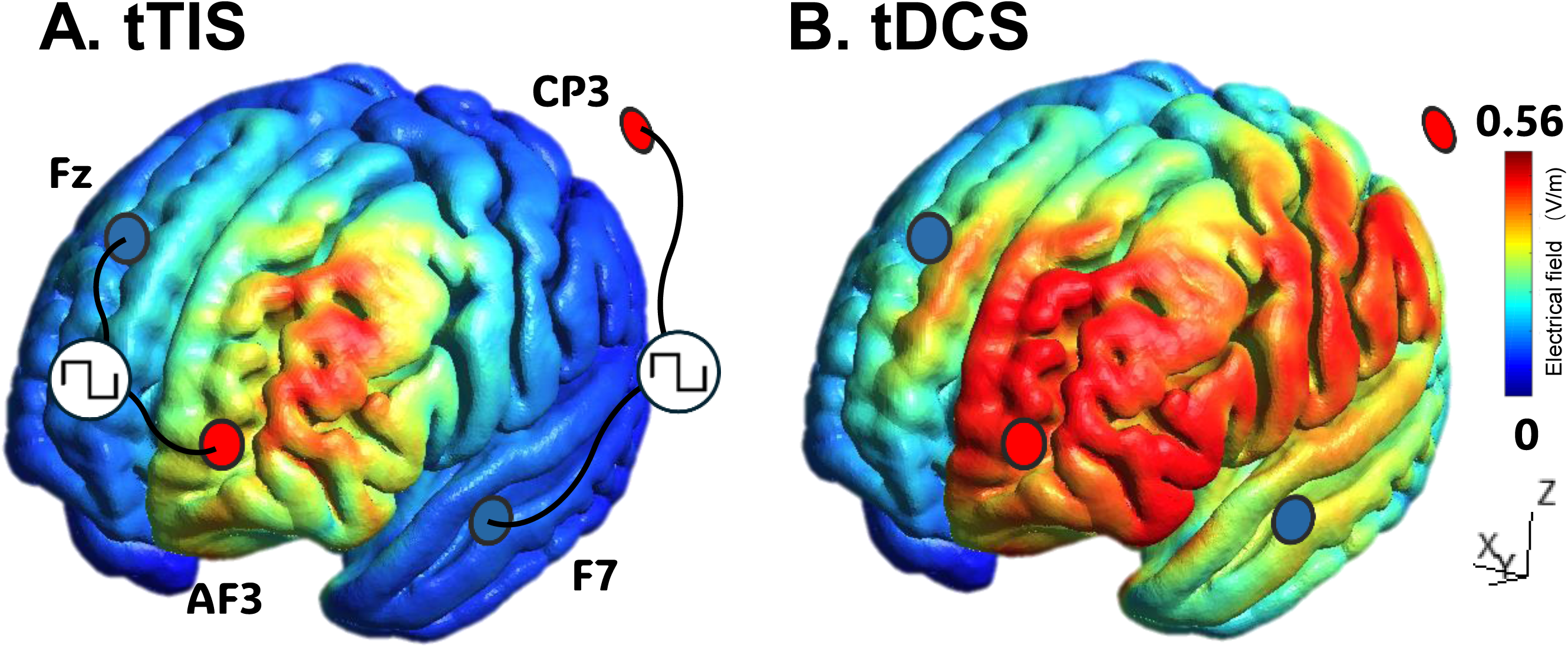
Electric-field simulations of the stimulation montage. (A) tTIS simulation using the experimental montage. The first pair of electrodes was placed at AF3 and Fz, while the second pair was placed at CP3 and F7. The injected current was set at 1.6 mA (peak to baseline) per electrode. (B) tDCS simulation using the same electrode placement and current intensity, shown for comparison.

### Statistical analysis

#### Trial-level analysis of prediction error

Single-trial FRN amplitudes were analyzed using linear mixed-effects models (LME; MATLAB fitlme) [26]. Behavioral prediction errors (PE) were coded as −1 (worse-than-expected), 0 (expected), and +1 (better-than-expected). The model included fixed effects of PE, Group (active vs. sham), Phase (pre vs. post), and their interactions: FRN amplitude ∼ PE × Group × Phase + (1 + PE | subject). If the maximal model failed to converge, a random-intercept model (1 | subject) was used. Fixed effects were evaluated using Wald z tests (two-tailed). Significant interactions were followed by simple-effects analyses across PE levels. Each session consisted of 150 trials. Behavioral negative prediction errors occurred relatively infrequently, yielding on average approximately 22 trials per participant per phase (range: 8–67).

#### Computational estimation of trial-level prediction errors

Given the limited number of trials for categorical negative prediction errors, model-derived prediction errors (PE_RW) were additionally estimated using a Rescorla–Wagner computational model [2] fitted to trial-wise behavioral data (Supplementary Section 3). This approach provided a continuous estimate of trial-wise prediction error and incorporated a larger proportion of trials (approximately 84 trials per phase). Single-trial FRN amplitudes were analyzed using an analogous LME model: FRN amplitude ∼ PE_RW × Group × Phase + (1 + PE_RW | subject). If the maximal model failed to converge, a simplified random-intercept model (1 | subject) was used. Follow-up analyses examined Group × Phase effects across PE_RW sign (PE_RW < 0, PE_RW = 0, PE_RW > 0).

#### ERP and behavioral analyses

Mean ERP and behavioral outcomes were analyzed using repeated-measures ANOVAs with Group (active vs. sham) as a between-subject factor and Phase (pre vs. post) as a within-subject factor. FRN amplitude was quantified as peak negativity within the 250–350 ms window at FCz, and SPN amplitude as mean voltage within the −200 to 0 ms window preceding feedback onset. Difference waves (no reward − reward) were used for visualization. For condition-level ERP analyses, negative prediction error trials were defined based on expectation–outcome mismatch (PE = −1). Greenhouse–Geisser corrections were applied where appropriate, with statistical significance set at *p* < .05.

### Power analysis and blinding

Sample size was determined a priori using G*Power (version 3.1) [27] for a repeated-measures ANOVA testing the Group × Phase interaction. Assuming a medium effect size (*f* = 0.25), *α* = .05, and power = 0.80, the required sample was 34 participants (17 per group).

### Integrity of the Blind

This study employed a single-blind design in which participants were unaware of the stimulation condition (active vs. sham). Due to technical constraints inherent to the tTIS setup, the experimenter administering stimulation was not blinded. To assess the integrity of blinding, participants were asked after each session to indicate whether they believed they had received active or sham stimulation. Blinding success was evaluated by comparing guessing accuracy against chance level (50%). Participants were unable to reliably identify the stimulation condition, with guessing accuracy not differing from chance, indicating preserved participant-level blinding.

## Results

### Demographic characteristics

Thirty-four participants were randomized to the active tTIS group (n = 17) or the sham stimulation group (n = 17). Demographic and clinical characteristics are summarized in Table S2. The two groups did not differ in Yale Food Addiction Scale scores, body mass index, age, education level, household income, or hunger ratings (all *p* > .05). Mild transient sensations (e.g., headache or tingling) were reported more frequently in the active group, but no participant reported lasting discomfort. Participants were unable to reliably identify stimulation condition (Table S3).

### Trial-level encoding of behavioral prediction errors

Single-trial FRN amplitudes scaled significantly with PE (*β* = 2.66, *SE* = 0.44, *z* = 5.99, *p* = 2.06 × 10⁻⁹), indicating robust trial-level sensitivity of FRN amplitudes to prediction errors during reward feedback. Significant PE × Phase (*β* = −1.69, *SE* = 0.61, *z* = −2.76, *p* = .006) and PE × Group interactions (*β* = −1.51, *SE* = 0.63, *z* = −2.39, *p* = .017) were observed. Importantly, a significant Group × Phase × PE interaction (*β* = 1.74, *SE* = 0.89, *z* = 1.97, *p* = .049) indicated that pre-to-post changes in the trial-level FRN–PE slope differed between the active and sham groups.

Figure 4 illustrates the trial-level relationship between FRN amplitude and prediction errors across groups and phases, based on fitted relationships from the mixed-effects model. Follow-up contrasts indicated that the Group × Phase effect was most pronounced for unfavorable outcomes (PE = −1; *β* = −3.45, *SE* = 0.97, *z* = −3.56, *p* = 3.70 × 10⁻⁴; Table S4). Subject-level FRN–PE slopes are provided in Supplementary Figure S1 for descriptive purposes.

**Figure 4.**
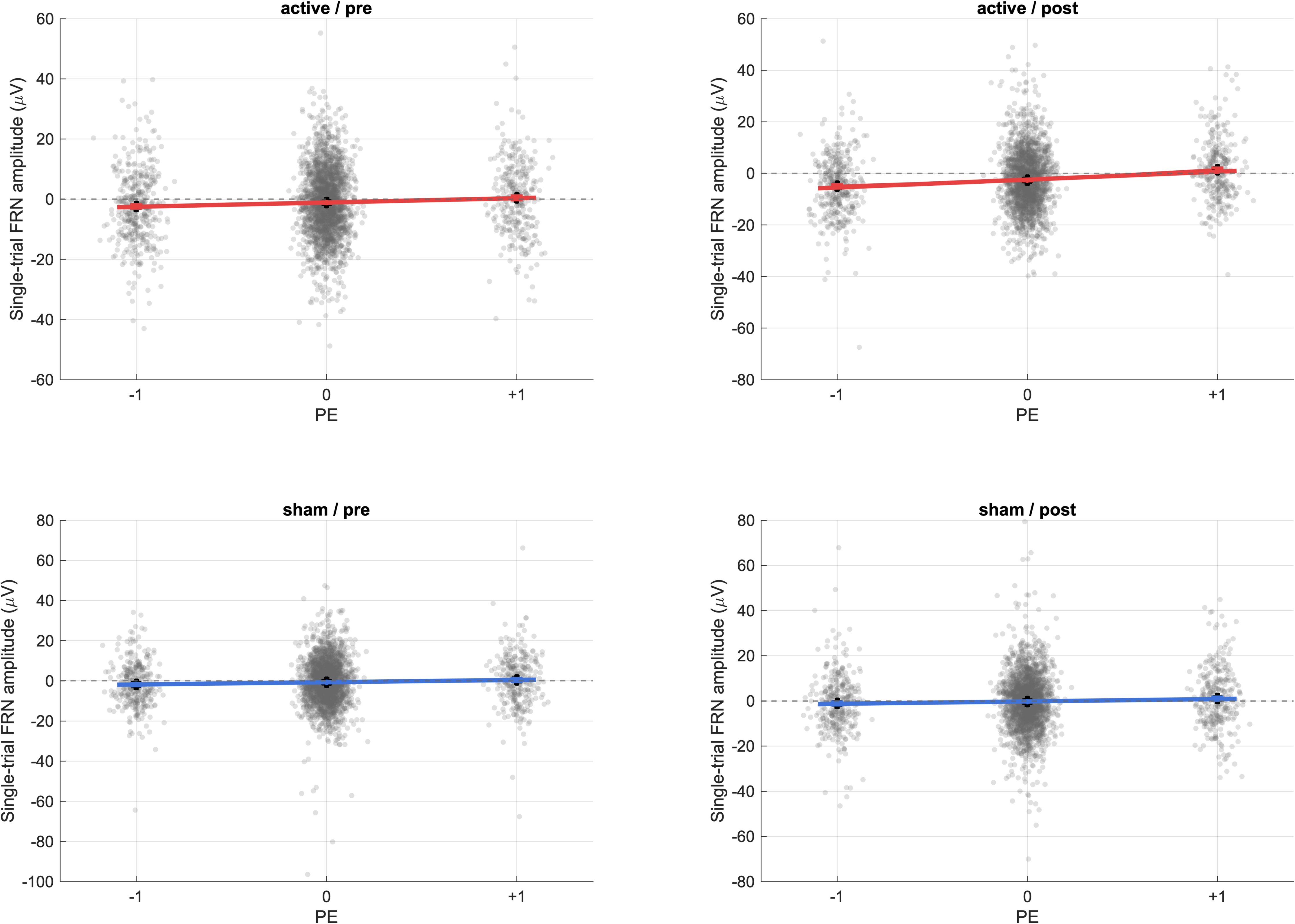
Trial-level FRN amplitudes as a function of behavioral prediction error category. Scatterplots show single-trial FRN amplitudes (µV) across behavioral prediction error categories (PE = −1, 0, +1), separately for the active and sham groups in the pre- and post-stimulation phases. Lines indicate fitted trial-level relationships shown for visualization.

### Trial-level encoding of model-based prediction errors

Single-trial FRN amplitudes scaled significantly with PE_RW (*β* = 2.94, *SE* = 0.45, *z* = 6.53, *p* = 6.62 × 10⁻¹¹), indicating robust trial-level sensitivity of FRN amplitudes to model-derived prediction errors. Significant PE_RW × Group (*β* = −1.42, *SE* = 0.64, *z* = −2.21, *p* = .027) and PE_RW × Phase interactions (*β* = −1.04, *SE* = 0.62, *z* = −1.69, *p* = .092) were observed. Critically, a significant Group × Phase × PE_RW interaction (*β* = 1.77, *SE* = 0.90, *z* = 1.98, *p* = .048) indicated differential pre-to-post changes in the trial-level FRN–PE_RW slope between groups.

Figure 5 illustrates the trial-level relationship between FRN amplitude and model-derived prediction errors across groups and phases, based on fitted relationships from the mixed-effects model. Follow-up contrasts indicated that the Group × Phase effect was most pronounced for negative prediction errors (PE_RW < 0; *β* = −2.46, *SE* = 0.56, *z* = −4.36, *p* = 1.29 × 10⁻⁵; Table S5). Subject-level slopes are provided in Supplementary Figure S2 for descriptive purposes.

**Figure 5.**
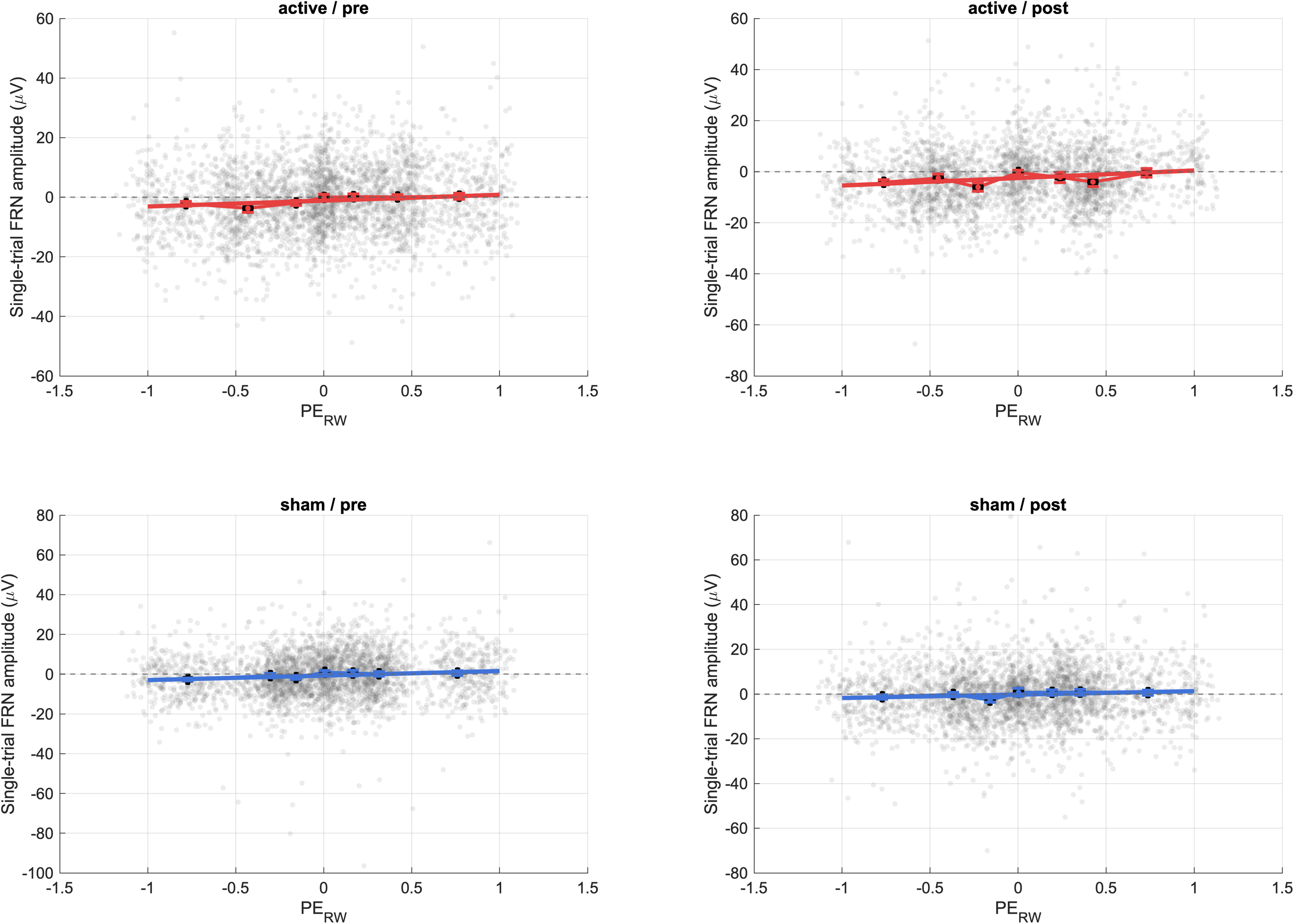
Trial-level FRN amplitudes as a function of model-derived prediction errors. Scatterplots show single-trial FRN amplitudes (µV) as a function of Rescorla–Wagner model-derived prediction errors (PE_RW), separately for the active and sham groups in the pre- and post-stimulation phases. Lines indicate fitted trial-level relationships shown for visualization.

### ERP indices of feedback and anticipation

#### FRN

To complement the trial-level mixed-effects analyses, condition-averaged FRN amplitudes were additionally examined for negative prediction errors (PE = −1; PE_RW < 0). This follow-up was guided by the trial-level localization analyses indicating that stimulation-related modulation was most pronounced for unfavorable outcomes (Tables S4–S5). These analyses yielded a consistent pattern of increased post-stimulation FRN negativity in the active group, with no corresponding change in the sham group (Figures S3–S4; Supplementary Section 6 for full statistics).

Figure 6 shows grand-average FRN waveforms and scalp topographies elicited by food reward outcomes in the active and sham groups before and after stimulation. FRN was observed as a fronto-central negative deflection peaking around 300 ms after feedback. Baseline FRN amplitudes did not differ between groups during the pre-stimulation phase (*t*_32_ = −0.35, *p* = .730, *d* = .12). A significant main effect of phase was observed (*F*_1,32_ = 6.13, *p* = .019, 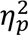 = .16), along with a trend-level Group × Phase interaction (*F*_1,32_ = 3.86, *p* = .058, 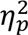 = .12). Follow-up analyses indicated that FRN amplitudes became significantly more negative after stimulation in the active group (*t*_16_ = 2.75, *p* = .014, *d* = .67), consistent with the trial-level findings, whereas no significant change was observed in the sham group (*t*_16_ = 0.44, *p* = .67, *d* = .11) (Figure 8A).

**Figure 6.**
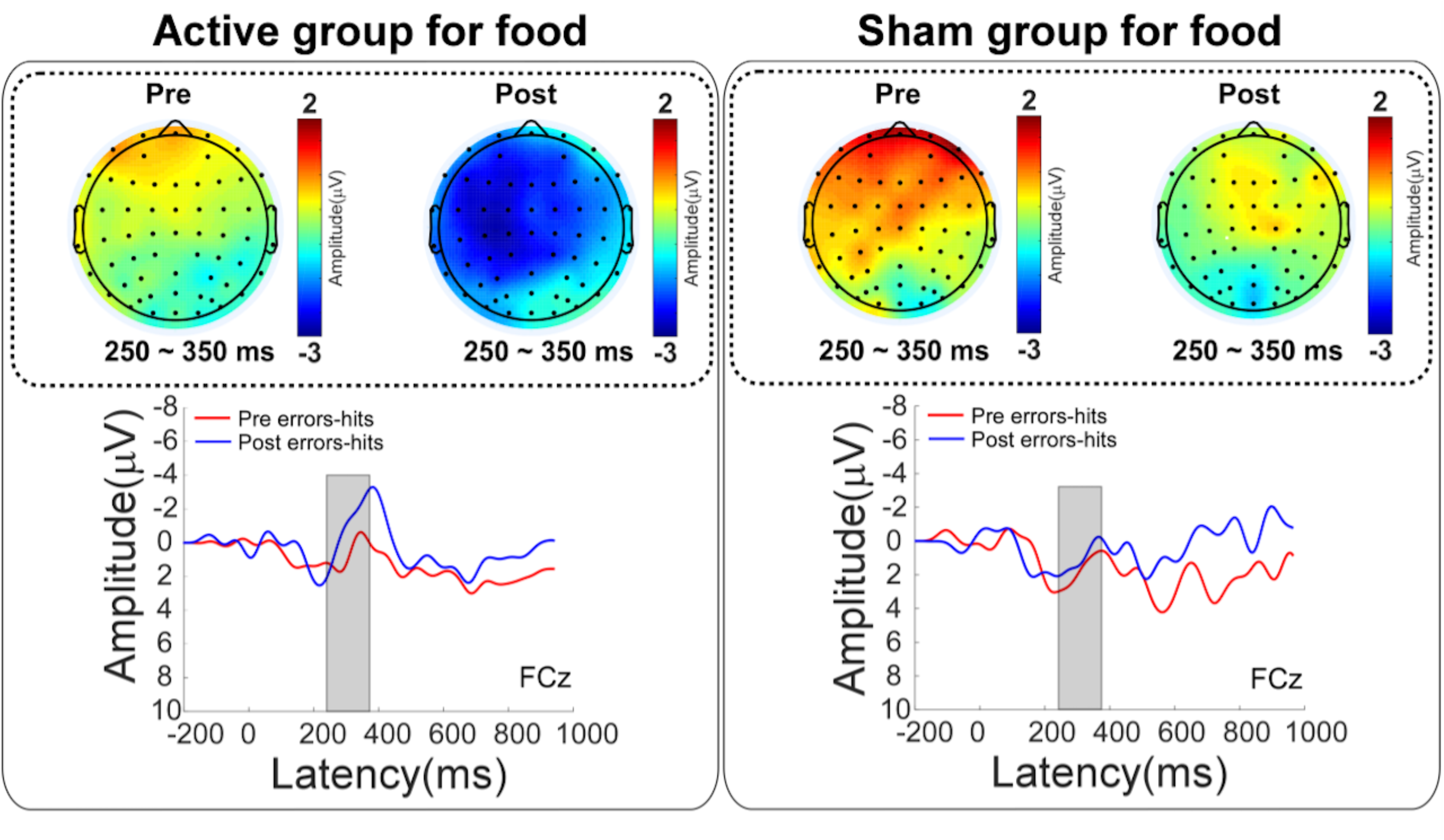
FRN elicited by food reward outcomes before and after stimulation. Grand-average ERP waveforms at FCz (top) and corresponding scalp topographies (bottom) are shown for the active tTIS group (left) and the sham group (right), separately for the pre- and post-stimulation phases. FRN was quantified as the peak negativity within the 250–350 ms time window following feedback onset (highlighted in gray). Waveforms are time-locked to feedback onset (0 ms) and plotted with pre-stimulation in red and post-stimulation in blue. Topographies depict mean amplitude within the FRN time window.

#### SPN

SPN waveforms and scalp distributions are shown in Figure 7. Baseline SPN amplitudes were comparable between groups (*t*_32_ = 0.60, *p* = .55, *d* = .21). No main effects of Group or Phase, nor a Group × Phase interaction, were observed (all *p* > .80). Quantitative SPN results are summarized in Figure 8B.

**Figure 7.**
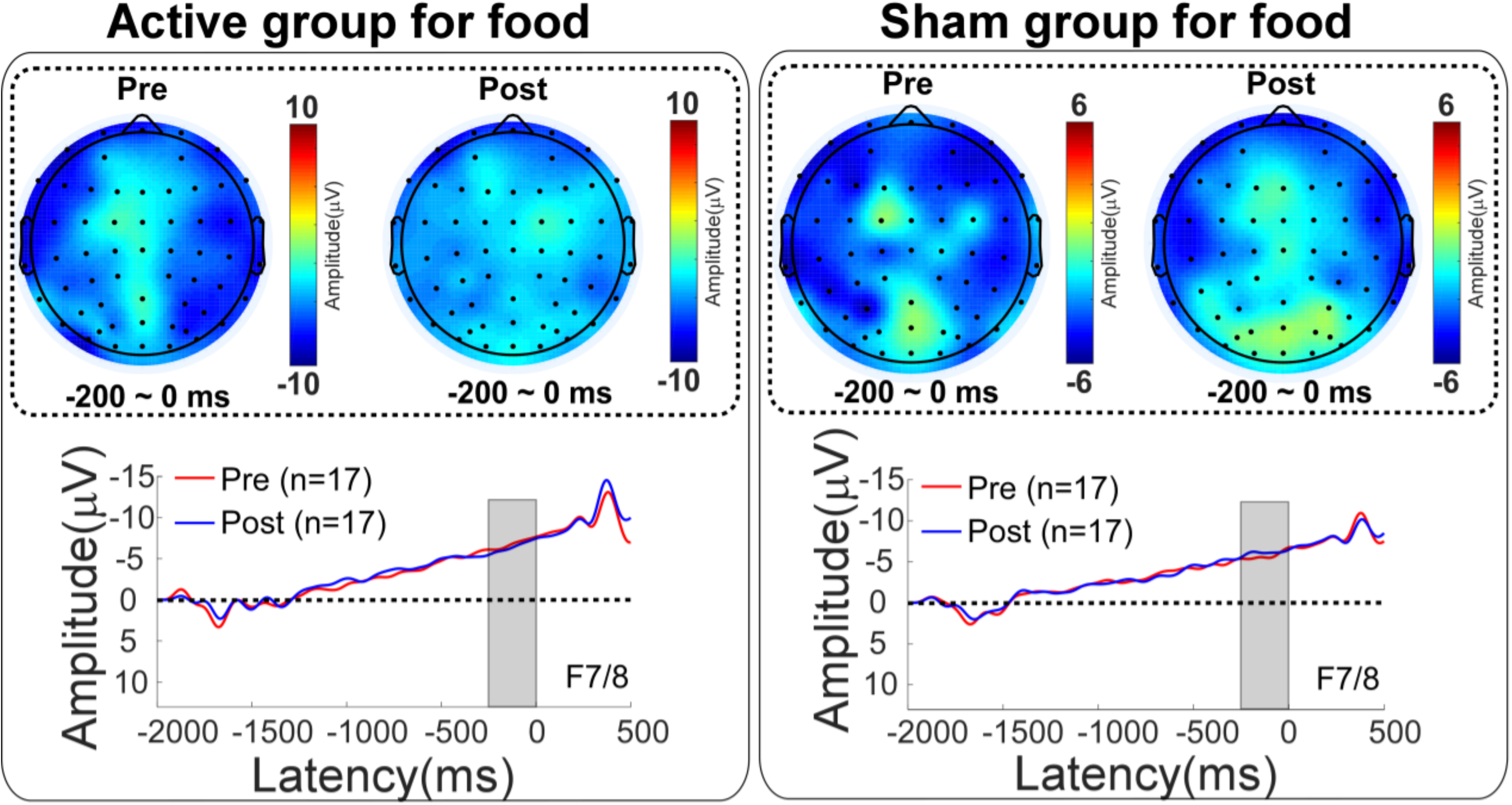
SPN during anticipation of food reward outcomes before and after stimulation. Grand-average SPN waveforms and scalp topographies are shown for the active tTIS group (left) and the sham group (right), separately for the pre- and post-stimulation phases. SPN was quantified as mean amplitude during the −200 to 0 ms interval prior to feedback onset (highlighted in gray). Waveforms are time-locked to feedback onset (0 ms) and plotted with pre-stimulation in red and post-stimulation in blue. Topographies depict mean amplitude within the SPN measurement window.

**Figure 8.**
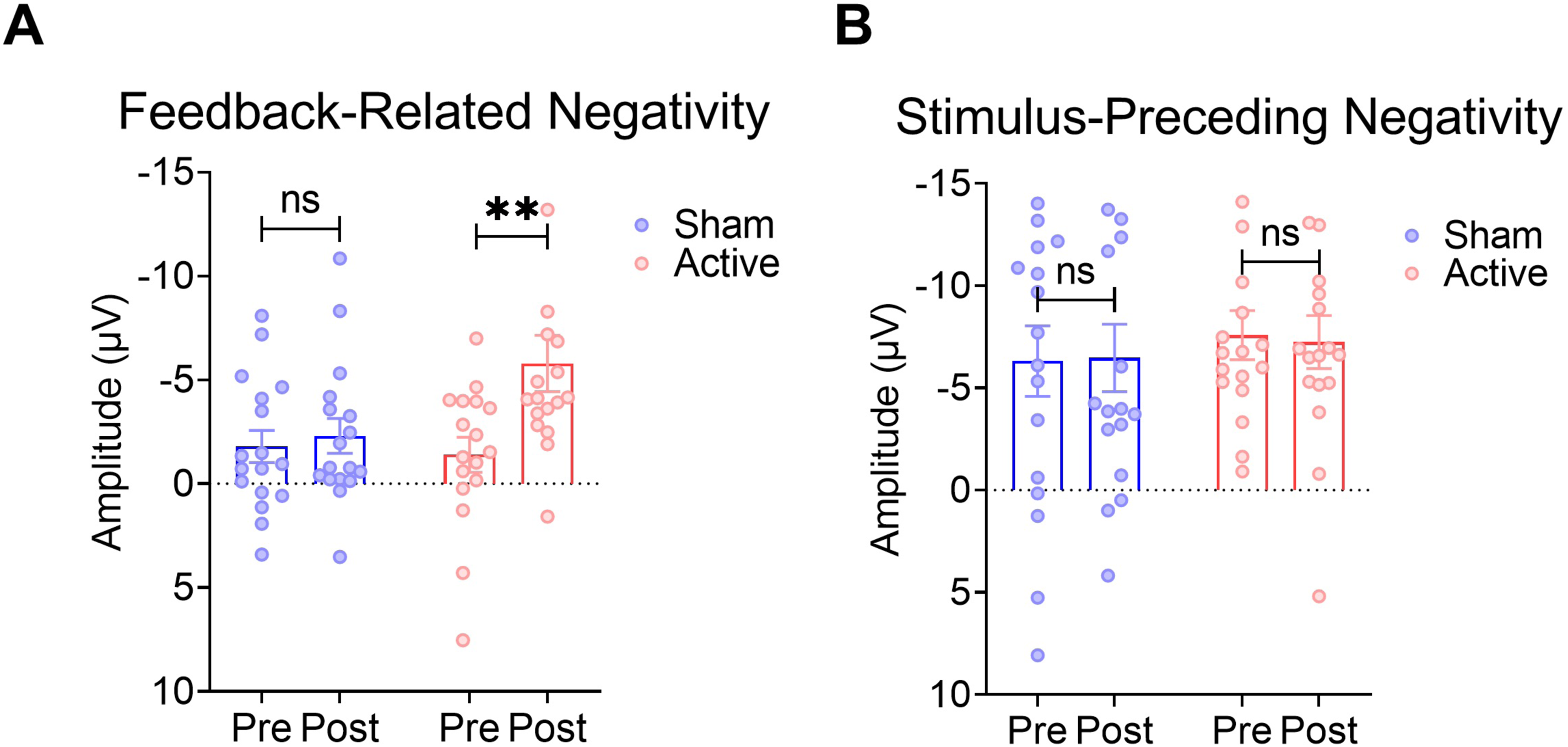
Quantitative ERP results for feedback evaluation and anticipation. (A) FRN amplitudes elicited by food reward feedback at FCz (250–350 ms), shown separately for the sham and active tTIS groups before (Pre) and after (Post) stimulation. (B) SPN amplitudes during the anticipatory period (−200 to 0 ms) prior to feedback onset. Bars represent group means (± SEM), and dots represent individual participants. Significance is denoted as *p* < .01 (**); *ns*, non-significant.

### Behavioral data

Behavioral performance during the incentive delay task was analyzed to examine whether stimulation-related neural effects were accompanied by behavioral changes. Reaction times showed a significant main effect of Phase (*F*_1,32_ = 10.73, *p* = .003, 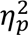 = .25), with faster responses post-stimulation, but no main effect of Group or Group × Phase interaction (both *p* > .05). Reward prediction rates did not differ as a function of Group, Phase, or their interaction for either high- or low-probability cues (all *p* > .05).

## Discussion

To our knowledge, the present study provides the first causal evidence that theta-band tTIS targeting the dACC modulates feedback-stage prediction error encoding during reward processing. In a randomized sham-controlled pre-post design, active stimulation altered the trial-level coupling between FRN amplitudes and prediction errors derived from both behavioral expectation-outcome mismatch and a Rescorla-Wagner computational model. Complementary ERP analyses converged with these findings, showing increased FRN negativity following active stimulation, whereas anticipatory activity indexed by SPN remained unchanged. Together, these results suggest that theta-band tTIS preferentially modulated the neural gain of prediction error encoding during feedback processing.

### Trial-level modulation of prediction error gain

The primary empirical contribution lies in the trial-level analyses linking FRN amplitudes to prediction errors. Single-trial FRN amplitudes scaled reliably with both categorical behavioral prediction errors and continuous prediction errors estimated from a Rescorla-Wagner computational model, indicating robust neural tracking of outcome discrepancies during feedback evaluation.

Critically, theta-band tTIS significantly modulated this relationship, as reflected by Group × Phase × prediction error interactions observed in the mixed-effects models. These results indicate that stimulation altered the strength of trial-level FRN–prediction error coupling across sessions, with differential pre-to-post changes between the active and sham groups. Follow-up contrasts further localized this modulation primarily to unfavorable outcomes (PE = −1; PE_RW < 0), consistent with previous findings showing stronger FRN responses to negative feedback during outcome evaluation [1,28].

Together, these findings suggest that theta-band tTIS altered the gain of prediction error encoding, as reflected in the sensitivity of FRN amplitudes to trial-by-trial outcome discrepancies, rather than producing a nonspecific shift in feedback-evoked activity. Subject-level slopes showed a similar numerical pattern but did not reach statistical significance. The primary inference therefore derives from the trial-level mixed-effects models, which leverage all available observations and provide greater statistical sensitivity to detect changes in this coupling.

### dACC theta modulation as a candidate mechanism

The present findings are consistent with the established role of the dACC in signaling outcome discrepancies and coordinating adaptive control through theta-band dynamics [7,29]. Converging electrophysiological and neuroimaging evidence indicates that medial frontal theta activity supports feedback evaluation, error monitoring, and adaptive control following unfavorable outcomes [7,30,31]. The FRN, a fronto-central negative deflection observed during outcome evaluation, has been closely linked to medial frontal theta activity and is widely interpreted as an electrophysiological index of prediction error processing [1,3,4].

Within this framework, the observed stimulation effects suggest that theta-band tTIS influenced the responsivity of dACC-centered evaluative circuits to deviations between expected and actual outcomes. Because the stimulation protocol generated a theta-frequency envelope targeting the dACC, the enhanced coupling between FRN amplitudes and prediction errors is consistent with the hypothesis that medial frontal theta dynamics regulate the neural gain of prediction error encoding.

However, the present EEG data do not allow direct differentiation between local modulation of dACC theta activity and indirect effects mediated through distributed cingulo-frontal networks. Given the dense anatomical and functional connectivity of the dACC, it is likely that stimulation influenced a broader feedback-evaluation network rather than a single isolated cortical node. Thus, the present findings are best interpreted as evidence that theta-band tTIS influenced prediction error encoding within a dACC-centered network, rather than indicating exclusive modulation of a single cortical site. Although electric-field modeling indicated peak envelope amplitude in medial frontal regions, the findings are best interpreted as network-level rather than strictly focal modulation. Future work combining tTIS with source-resolved EEG, connectivity analyses, or multimodal imaging will be important for clarifying the network-level mechanisms underlying these effects.

### Dissociation between feedback evaluation and anticipatory processing

Theta-band tTIS exerted stage-specific effects on reward processing. Whereas stimulation reliably altered feedback-stage FRN–prediction error coupling, anticipatory activity indexed by SPN remained unchanged. Because SPN has been associated with anticipatory attention, motivational engagement, and preparatory processes occurring prior to feedback delivery [32–34], this dissociation suggests that theta-band modulation of dACC-centered networks preferentially influences evaluative computations engaged during outcome processing rather than anticipatory preparatory processes [1,3,5,35].

This stage specificity also aligns with previous observations of an imbalanced anticipatory–feedback profile in food addiction under satiety, characterized by relatively elevated anticipatory signals but reduced feedback-based updating relative to healthy controls [18]. Within this context, the present findings suggest that a single session of theta-band tTIS may acutely modulate feedback updating mechanisms that support prediction error signaling, without substantially altering anticipatory expectancy signals. Repeated stimulation or alternative stimulation parameters may be required to influence anticipatory components such as SPN.

### Implications for non-invasive modulation of deep reward-monitoring circuits

tTIS has emerged as a promising non-invasive approach for modulating deep cortical structures that are difficult to access with conventional stimulation techniques [14,15]. Although early studies primarily relied on computational modeling or indirect behavioral outcomes, recent human work has begun to provide evidence for functional modulation of targeted circuits [16,17]. By demonstrating that theta-band tTIS altered trial-level FRN–prediction error coupling, the present study extends this literature beyond mean-amplitude ERP effects and illustrates how tTIS can be used to causally probe prediction-error-related computations in deep medial frontal circuits.

More broadly, these findings support the use of tTIS as a mechanistic tool for interrogating deep reward-monitoring circuits in humans. They also motivate future work to test whether repeated stimulation can produce more sustained changes in prediction error signaling, downstream behavioral adaptation, and potentially clinically relevant reward-learning phenotypes.

### Limitations and future directions

Several limitations should be acknowledged. First, although the sample size was sufficient to detect the primary interaction effects and trial-level mixed-effects models improved sensitivity by leveraging all available observations, statistical power may have been limited for detecting subtler between-group differences in subject-level summary indices such as individual FRN–prediction error slopes [26]. Second, only short-term effects of a single stimulation session were assessed; future studies should examine whether repeated stimulation produces sustained changes in prediction error sensitivity and related behavioral adaptation. Third, although the stimulation target was defined in the dACC, the present EEG data do not permit precise source-level attribution, and stimulation may have influenced a broader cingulo-frontal network. Finally, reward processing depends on distributed circuits beyond the dACC, including striatal and prefrontal systems [36]; combining tTIS with multimodal imaging or connectivity analyses may help clarify the network-level mechanisms underlying the present effects. Although mild sensations were more frequently reported in the active condition, participants were unable to reliably identify stimulation condition, suggesting that expectancy-related confounds are unlikely to account for the observed effects.

### Conclusion

Theta-band tTIS targeting the dACC modulated feedback-stage prediction error encoding during reward processing in participants from the general population with elevated food addiction-related symptoms. Trial-level mixed-effects analyses showed altered FRN–prediction error coupling for both behavioral and model-derived prediction errors, with effects localized primarily to unfavorable outcomes. Complementary ERP analyses converged with these findings, showing enhanced FRN negativity after active stimulation without changes in SPN. Given that prediction error encoding reflects a domain-general computational process, these findings are consistent with a domain-general role of prediction error encoding in outcome evaluation. Together, these results demonstrate that medial frontal theta dynamics contribute to the neural gain of prediction error encoding and support tTIS as a promising tool for probing deep, frequency-specific computations in dACC-centered reward-monitoring circuits.

## Acknowledgments

This work was supported by the Chinese National Programs for Brain Science and Brain-like Intelligence Technology (Grant No. 2021ZD0202104) and the International Cooperation and Exchange Program of the National Natural Science Foundation of China (Grant No. 32161133004).

## CRediT authorship contribution statement

**Yatong Wen:** Conceptualization, Methodology, Investigation, Formal analysis, Data curation, Visualization, Writing – original draft. **Yonghui Li:** Funding acquisition, Supervision, Writing – review & editing. **Yiheng Tu:** Funding acquisition, Supervision, Writing – review & editing. **Mo Wang:** Methodology, Software. **Xinwen Dong:** Validation, Writing – review & editing. **Mateusz Gola:** Writing – review & editing. **Quanying Liu:** Methodology.

## Declaration of competing interest

The authors declare that they have no known competing financial interests or personal relationships that could have appeared to influence the work reported in this paper.

## Preprint

A preprint version of this manuscript has been deposited on the bioRxiv preprint server and is disclosed in accordance with journal policy. DOI: 10.64898/2025.12.20.695275.

## Data availability

De-identified behavioral data and derived ERP measures supporting the findings of this study are available at OSF: https://doi.org/10.17605/OSF.IO/N28A6. Analysis scripts will be made publicly available upon publication.

## Declaration of generative AI and AI-assisted technologies in the manuscript preparation process

During the preparation of this manuscript, the authors used ChatGPT (OpenAI) for language editing and improving readability. The authors reviewed and edited all content to ensure accuracy and take full responsibility for the final manuscript.

---

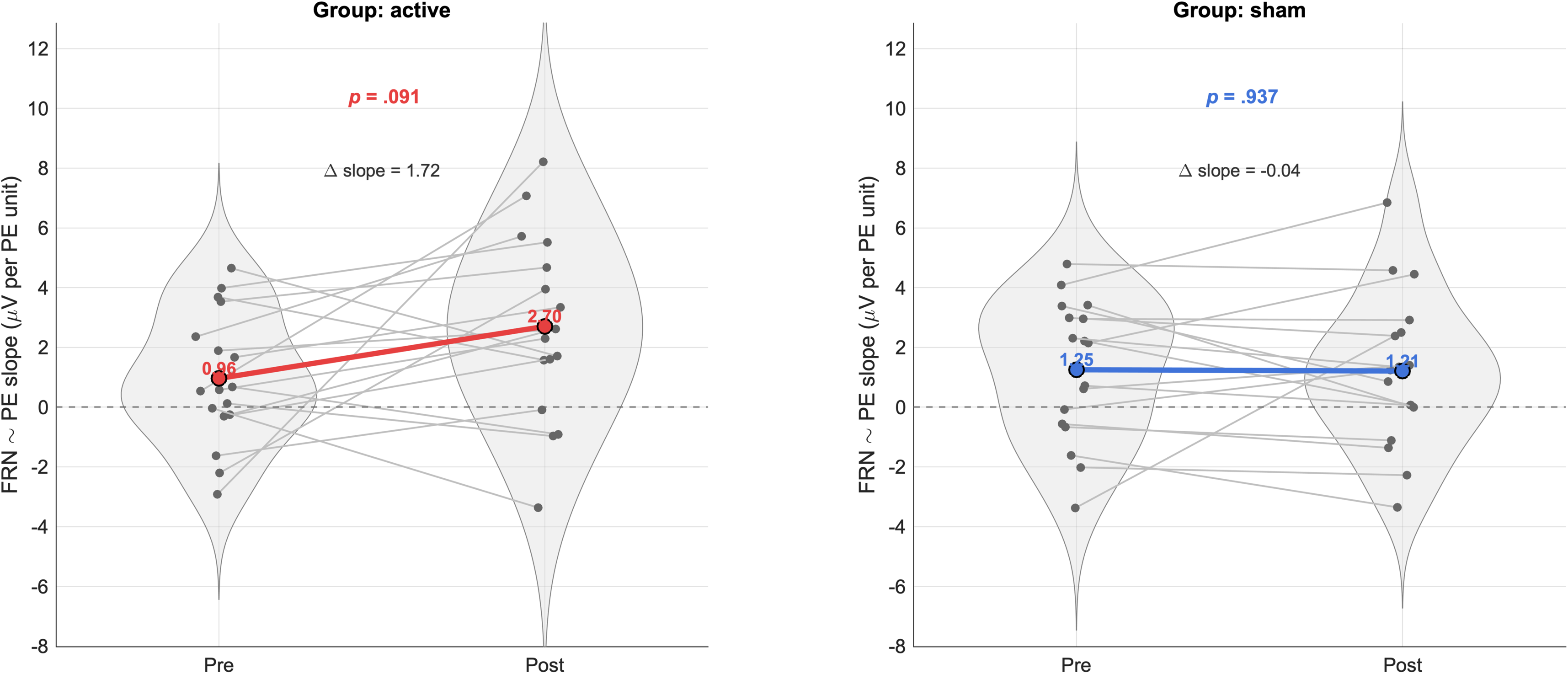
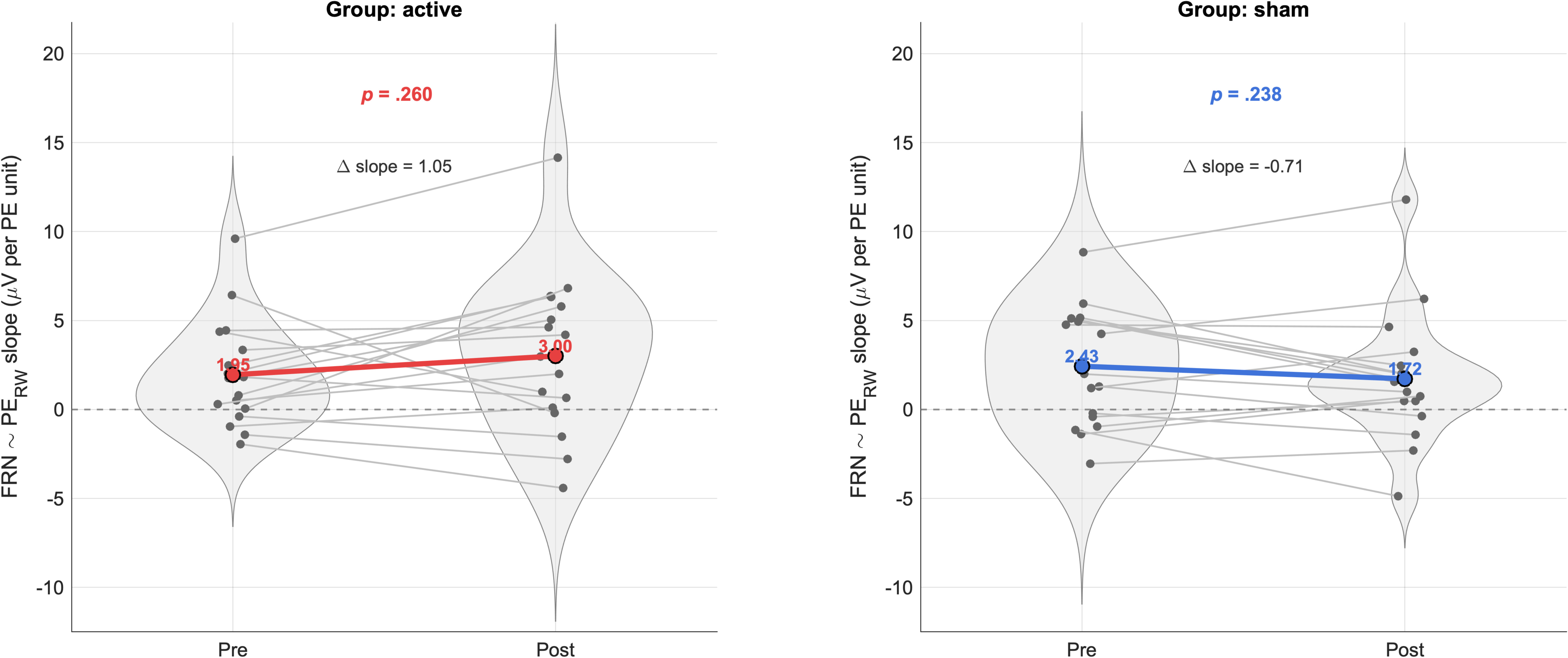
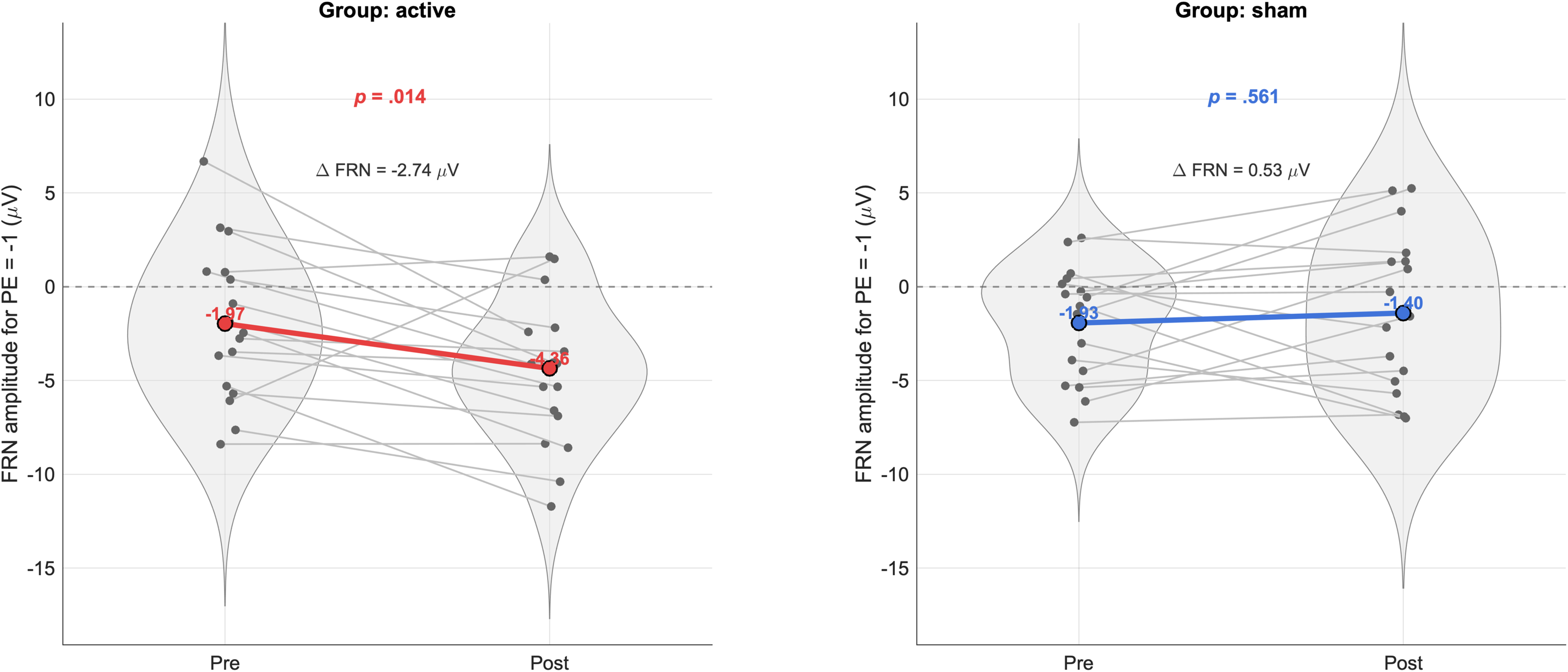
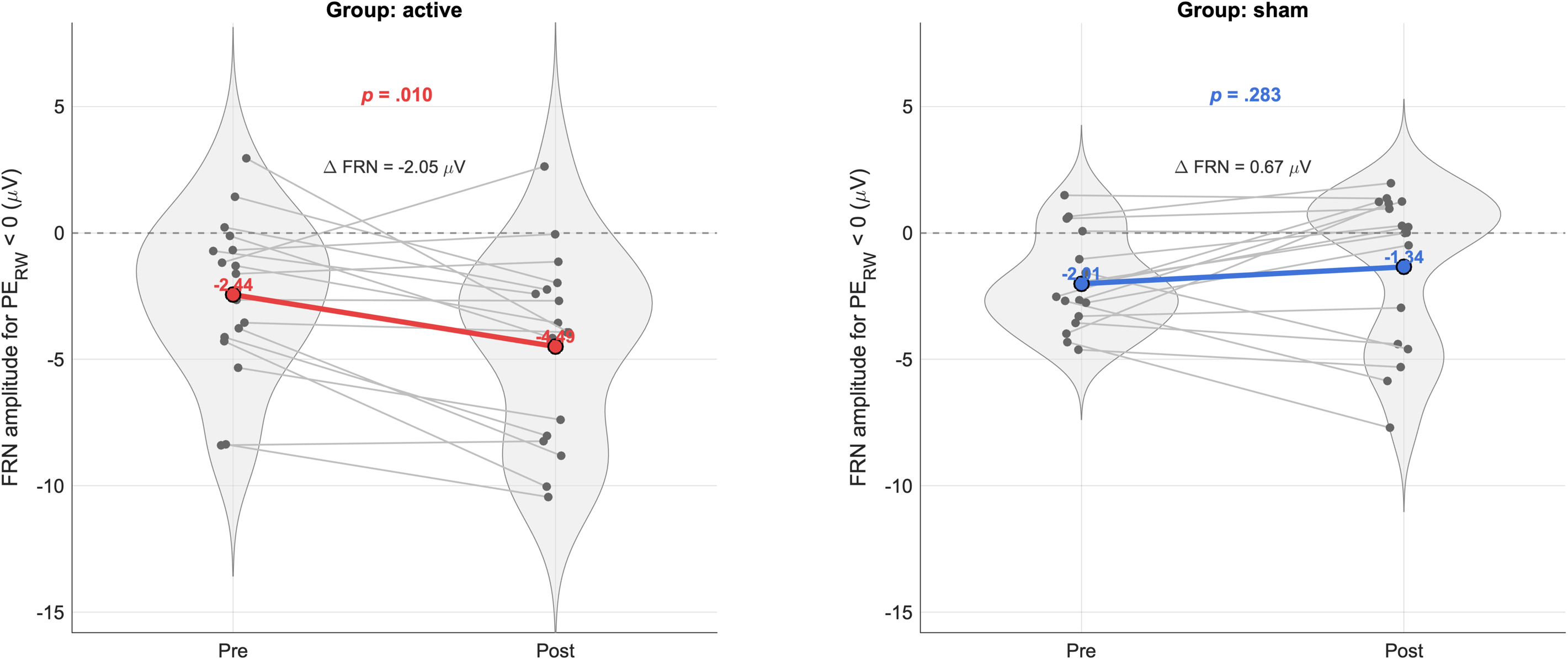

## Notes

### Competing Interest Statement

The authors have declared no competing interest.

### Summary of Updates

This version of the manuscript has been substantially revised. Specifically, the title, abstract, and introduction have been updated to improve clarity and conceptual framing. The organization of the Results section has been restructured to emphasize trial-level analyses and their primary role in supporting the main conclusions. The Discussion has been revised to better align with the updated analytical framework and interpretation. References have been updated accordingly.

