## Supplementary Materials for "Theta-Band Temporal Interference Stimulation Targeting the dACC Modulates Neural Gain of Prediction Error Encoding"

**Supplementary Section 1. Reward image matching**

Fifteen independent participants (mean age = 23.19 years, SD = 3.33), who did not take part in the main experiment, rated 124 candidate food images using an online questionnaire. Images were evaluated on arousal, pleasantness, and familiarity using 10-point Likert scales.

Based on these ratings, 75 food reward images were selected for use in the incentive delay task. Selection criteria required arousal and familiarity ratings of at least 5, while pleasantness ratings were distributed across low, medium, and high levels (25 images per category) to ensure variability in subjective reward value.

All food reward images (720 × 540 pixels) depicted the target item on a black background. Non-reward control images were generated by segmenting and recombining food reward images into mosaic patterns using MATLAB, with brightness gradients applied to match low-level visual properties. Descriptive statistics for the selected food reward images are provided in Table S1, demonstrating adequate arousal, pleasantness, and familiarity across the stimulus set (all $p > .05$).

**Table S1. Reward image matching**

| Reward | Pleasantness (M ± SD) | Arousal (M ± SD) | Familiarity (M ± SD) | *t* | *p* |
|---|---|---|---|---|---|
| Food (N = 75) | 6.63 ± 0.69 | 6.38 ± 0.73 | 7.73 ± 0.75 | 1.73 | .088 |

***Note.*** *M represents the mean and SD represents the standard deviation.*

**Supplementary Section 2. tTIS simulation**

We employed the SimNIBS software package to perform finite element method (FEM) simulations [1] (available for download at *https://simnibs.github.io/simnibs/build/html/installation/simnibs_installer.html*). With MNI152 template, we used the CHARM pipeline integrated within SimNIBS to process T1-weighted image and generate FEM model [2]. The input image is segmented into different brain tissues and then converted into meshes. Each mesh is assigned a specific conductivity value based on the tissue type. For our simulations, we adopted the default conductivity settings: White Matter (0.126 S/m), Gray Matter (0.275 S/m), CSF (1.654 S/m), Compact Bone (0.008 S/m), Spongy Bone (0.025 S/m), Scalp (0.465 S/m), and Eyeballs (0.5 S/m).

The FEM model assumes there are no internal current sources or sinks within the brain, meaning no internal electrical activity occurs within the elements. This allows for the calculation of the electric field in each mesh using Poisson's equation：

$$\nabla\cdot(\sigma E) = -\nabla\cdot(\sigma\nabla V) = 0,$$

Here, the electric field (E) is calculated based on tissue conductivity (σ) and potential distribution (V). The FEM approach is used to assemble a global matrix, and boundary conditions are defined based on the electrode settings consistent with the stimulator configuration. The potential from the electrode (V0) is determined by factors such as the material, size, and positioning of the electrode, and the Dirichlet boundary condition is applied to ensure the electrode potential is accurately represented.

$$V = V_0,$$

After assembling the global matrix and applying the boundary conditions, the system is solved numerically to obtain the distribution of the electric field with one pair electrode.

The lead field matrix is then calculated by considering all candidate electrodes based on the EEG-10/10 system. This matrix defines how each electrode contributes to the electric field distribution across different regions of the brain. Specifically, the lead field matrix is derived by calculating the electric field components along the x, y, and z axes for each electrode configuration. The lead field matrix enables us to predict the electric field generated at any location in the brain in response to stimulation, represented as:

$$E = Ls,$$

where E is the electric field, L is the lead field matrix, and s represents the injected currents. Subsequently, we determined the envelope of the electric field distribution using the following equation [3,4] :

$$|\vec{E}|_{max} = 2\max_{\alpha}\min\left(||\vec{E}_1|||\cos\alpha|, ||\vec{E}_2|||\cos(\alpha-\phi)|\right)$$

where $E_1$ and $E_2$ are the first and second distinct fields, $\phi$ is the angle spanned by $E_1$ and $E_2$, and $\alpha$ is the angle between $E_1$ and the projection of $n$ on that plane. 1D-search over $\alpha \in [0, 2\pi)$ can be performed to find the maximal modulation where $E_1$, $E_2$ and $\phi$ are known from computational model.

Once this relationship between the electric field and the currents is established, we can optimize the electrode configuration to achieve optimal stimulation. In this study, we employed the MOVEA, a multi-objective optimization algorithm that simultaneously optimizes the trade-offs between intensity and focality [5]. Here, intensity refers to the electric field strength at the target areas, while focality describes the mean of the field across the entire brain. We selected optimal configurations from the Pareto front calculated by the algorithm (see Figure 3). The electrode placements for targeting the dorsal anterior cingulate cortex (dACC) were determined as follows: the first electrode pair was positioned at AF3 (positive) and Fz (negative), while the second pair was placed at CP3 (positive) and F7 (negative). The MNI-to-Talairach coordinate conversion was performed using an online tool available at: https://bioimagesuiteweb.github.io/bisweb-manual/tools/mni2tal.html.

**Supplementary Section 3: Computational estimation of trial-level prediction errors**

To obtain a continuous estimate of trial-level prediction errors during reward learning, a Rescorla–Wagner–type update rule was applied to participants' single-trial behavioral data from the incentive delay task.

*Model specification*

For each participant, expected value $V_t$ was updated on each trial according to the standard Rescorla–Wagner learning rule:

$$V_{t+1} = V_t + \alpha \times \delta_t$$

Where $\delta_t$ represents the prediction error on trial $t$, defined as:

$$\delta_t = R_t - V_t$$

And $R_t$ denotes the outcome of the trial (reward = 1, no reward = 0). The learning rate $\alpha$ ($0 < \alpha < 1$) controls the degree to which new outcomes update the expected value.

*Model fitting procedure*

The learning rate parameter $\alpha$ was estimated separately for each participant using maximum likelihood estimation based on trial-level behavioral data. Model fitting was implemented using custom MATLAB scripts, with parameter optimization constrained within plausible bounds to ensure numerical stability. The primary aim of this procedure was to obtain reliable trial-level estimates of prediction errors rather than to fully characterize individual learning strategies.

*Derivation of trial-level prediction errors*

Once fitted, the model was used to generate trial-level prediction errors ($\delta_t$) for each participant and phase (pre- and post-stimulation). These continuous prediction error values were then aligned with corresponding EEG trials and entered into subsequent analyses relating FRN amplitudes to model-derived prediction errors.

*Statistical linkage to EEG data*

Trial-level FRN amplitudes were related to model-derived prediction errors using linear mixed-effects models, with FRN amplitude as the dependent variable, prediction error as a fixed effect, and subject as a random intercept. Group (active tTIS vs. sham) and phase (pre vs. post) were included as additional fixed effects, allowing assessment of how stimulation modulated the relationship between FRN and internal prediction error signals. In addition, subject-level FRN–prediction error slopes were estimated separately for each participant and phase to provide complementary descriptive indices of individual FRN–prediction error coupling.

It should be noted that the RW model was used here as a minimal and well-established framework to capture trial-level prediction errors, rather than to provide a comprehensive account of individual learning strategies.

**Supplementary Section 4. Demographic and reported sensation after tTIS**

*Demographic characteristics*

Demographic and clinical characteristics of participants in the active tTIS and sham groups are summarized in Table S2. The two groups did not differ significantly in age, BMI, YFAS scores, hunger level, education status, or household income (all $p > .05$).

Self-reported sensations during stimulation are summarized in **Table S3**. Mild sensations (e.g., tingling, scalp discomfort) were more frequently reported in the active tTIS group, whereas no participant reported lasting discomfort or was able to identify group assignment.

**Table S2. Demographic and clinical characteristics of participants in both the active tTIS and sham stimulation groups**

| Variable | TI Active Group (N = 17) | Sham stimulation Group (N= 17) |
|---|---|---|
| Age | 26.29 (SD = 5.84) | 24.35 (SD = 2.98) |
| Student Status | 13 | 13 |
| Only Child | Yes: 10 | Yes: 9 |
| Monthly Household Income (CNY) | > 10,000：7, <3000：1 | > 10,000：8, <3000：2 |
| Medical Conditions | Hypertension: 1 Dyslipidemia: 1 Other: 1 | Hypertension: 1 Dyslipidemia: 1 Other: 1 |
| BMI | 27.66 (SD = 4.52) | 27.62 (SD = 5.24) |
| YFAS | 6.00 (SD = 1.06) | 5.41 (SD = 1.00) |
| Hunger Level | 1.71(SD = 0.77) | 2.24 (SD = 0.90) |

***Note.*** *$*p < .05$, $**p < .01$, $***p < .001$*

**Table S3. The prevalence of reported sensation after active TI and sham stimulation groups**

| Sensation | TI Active Group (N = 17) | Sham stimulation Group (N= 17) |
|---|---|---|
| Headache | 47.1% (None), 35.3% (Mild), 17.6% (Moderate) | 64.7% (None), 23.5% (Mild), 11.8% (Moderate) |
| Neck Pain | 82.4% (None), 17.6% (Mild) | 94.1% (None), 5.9% (Mild) |
| Scalp Pain | 23.5% (None), 52.9% (Mild), 23.5% (Moderate) | 35.3% (None), 35.3% (Mild), 29.4% (Moderate) |

| | | |
|---|---|---|
| Scalp Injury | 70.6% (None), 29.4% (Mild) | 76.5% (None), 23.5% (Mild) |
| Tingling | 11.8% (None), 64.7% (Mild), 23.5% (Moderate) | 58.8% (None), 23.5% (Mild), 17.7% (Moderate) |
| Skin Redness | 82.4% (None), 17.6% (Mild) | 88.2% (None), 11.8% (Mild) |
| Drowsiness | 41.2% (None), 35.3% (Mild), 23.5% (Moderate) | 47.1% (None), 23.5% (Mild), 29.4% (Moderate) |
| Difficulty Concentrating | 47.1% (None), 52.9% (Mild) | 76.5% (None), 23.5% (Mild) |
| Acute Mood Changes | 88.2% (None), 11.8% (Mild) | 82.4% (None), 17.6% (Mild) |

***Note.*** *Percentages indicate the proportion of participants reporting each sensation level.*

### Supplementary Section 5. Follow-up analyses of FRN–PE interactions

To localize the significant Group × Phase × PE interaction, we conducted follow-up simple-effects analyses testing the Group × Phase contrast separately at each PE level (PE = −1, 0, +1). Results are summarized in Table S4. In addition, subject-level FRN–PE slopes are shown in Figure S1 as descriptive indices of individual FRN–PE slopes. Although slopes showed a numerically larger pre-to-post increase in the active group, these differences did not reach statistical significance (Δ slope = 1.72, $p$ = .091, $d$ = .45), with no apparent change in the sham group (Δ slope = −0.04, $p$ = .937, $d$ = .02).

**Table S4. Follow-up simple-effects analyses of the Group × Phase contrast at each PE level**

| PE | $\beta$ | $SE$ | $z$ | $p$ | 95% CI |
|---|---|---|---|---|---|
| −1 | −3.45 | 0.97 | −3.56 | $3.70 \times 10^{-4}$ | [−5.35, −1.55] |
| 0 | −1.71 | 0.43 | −3.94 | $8.07 \times 10^{-5}$ | [−2.56, −0.86] |
| 1 | 0.03 | 1 | 0.03 | .974 | [−1.94, 2.00] |

***Note.*** *Follow-up simple-effects analyses were conducted to decompose the significant Group × Phase × PE interaction from the LME model predicting single-trial FRN amplitudes. For each PE level, the Group × Phase contrast tests whether the pre-to-post change in FRN differs between the active and sham groups (difference-in-*

*differences). p values are based on normal approximation (two-tailed).*

To further characterize the significant Group × Phase × PE_RW interaction, we conducted follow-up simple-effects analyses testing the Group × Phase contrast separately within three PE_RW bins (PE_RW<0, PE_RW=0, PE_RW>0). Results are summarized in Table S5. In addition, subject-level FRN–PE_RW slopes are shown in Figure S2 for descriptive visualization. Although slopes showed a numerically increasing trend from pre to post in the active group (Δ slope = 1.05, $p$ = .260, $d$ = .29) and a decreasing trend in the sham group (Δ slope = −0.71, $p$ = .238, $d$ = .30), these differences did not reach statistical significance.

**Table S5. Follow-up simple-effects analyses of the Group × Phase contrast across PE_RW bins**

| PE_RW_bin | *β* | *SE* | *z* | *p* | 95% CI |
|---|---|---|---|---|---|
| PE_RW<0 | −2.46 | 0.56 | −4.36 | $1.29 \times 10^{-5}$ | [−3.56, −1.35] |
| PE_RW=0 | −1.74 | 0.43 | −4.03 | $5.67 \times 10^{-5}$ | [−2.59, −0.89] |
| PE_RW>0 | −1.05 | 0.56 | −1.89 | .059 | [−2.14, 0.04] |

***Note.*** *Follow-up simple-effects analyses were conducted to further characterize the significant Group × Phase × PE_RW interaction from the LME model predicting single-trial FRN amplitudes. For each PE_RW bin, the Group × Phase contrast tests whether the pre-to-post change in FRN differs between the active and sham groups (difference-in-differences). p values are based on normal approximation (two-tailed).*

### Supplementary Section 6. Condition-averaged FRN responses to negative prediction errors

To complement the trial-level mixed-effects analyses, we examined condition-averaged FRN amplitudes for negative prediction errors at the subject level. Specifically, FRN amplitudes were averaged across trials with negative behavioral prediction errors (PE = −1) and across trials with negative model-derived prediction errors (PE_RW < 0), separately for each participant and phase (pre vs. post). Pre-to-post changes were assessed using paired-samples t tests within each group.

Averaging FRN amplitudes across trials with negative behavioral prediction errors (PE = −1) revealed a significant post-stimulation increase in FRN negativity in the active group (mean_pre = −1.62 μV, mean_post = −4.36 μV, ΔFRN = −2.74 μV, $t_{16}$ = 2.78, $p$ = .014, $d$ = .69), whereas no significant change was observed in the sham group (mean_pre = −1.93 μV, mean_post = −1.40 μV, ΔFRN = 0.53 μV, $t_{16} = -0.59$, $p = .561$, $d = .14$) (Figure S3).

Averaging FRN amplitudes across trials with negative model-derived prediction errors (PE_RW < 0) yielded a comparable pattern, with a post-stimulation increase in FRN negativity in the active group (mean_pre = −2.26 μV, mean_post = −4.31 μV, ΔFRN = −2.05 μV, $t_{16}$ = 2.97, $p = .010$, $d = .74$), and no corresponding change in the sham group (mean_pre = −2.01 μV, mean_post = −1.34 μV, ΔFRN = 0.67 μV, $t_{16} = -1.11$, $p = .283$, $d = .27$) (Figure S4).

**Supplementary Figure Legends**

**Supplementary Figure S1. Subject-level slopes summarizing trial-level FRN–PE coupling.** Subject-level regression slopes (μV per PE unit) summarizing trial-level coupling between single-trial FRN amplitudes and categorical behavioral prediction errors (PE) are shown separately for the active theta-band tTIS group (left) and the sham group (right), in the pre- and post-stimulation phases. Each dot represents one participant, with paired lines connecting pre and post slopes. Violin plots depict the distribution of individual slopes within each phase.

**Supplementary Figure S2. Subject-level slopes summarizing trial-level FRN–PE_RW coupling.** Subject-level regression slopes (μV per PE unit) summarizing trial-level coupling between single-trial FRN amplitudes and model-derived prediction errors (PE_RW) are shown separately for the active theta-band tTIS group (left) and the sham group (right), in the pre- and post-stimulation phases. Each dot represents one participant, with paired lines connecting pre and post slopes. Violin plots depict the distribution of individual slopes within each phase.

**Supplementary Figure S3. Mean FRN for negative behavioral prediction errors (PE = −1).** Condition-averaged feedback-related negativity (FRN) amplitudes for unfavorable outcomes (negative behavioral prediction errors; PE = −1) during food reward processing, shown separately for the active theta-band tTIS and sham groups before and after stimulation. This mean-level follow-up was guided by trial-level localization results showing that stimulation-related modulation was most pronounced for negative prediction errors (Table S4). FRN amplitudes became more negative after stimulation in the active group, whereas no corresponding change was observed in the sham group.

**Supplementary Figure S4. Mean FRN for negative model-based prediction errors (PE_RW < 0).** Condition-averaged feedback-related negativity (FRN) amplitudes for unfavorable outcomes (negative model-derived prediction errors; PE_RW < 0) during food reward processing, shown separately for the active theta-band tTIS and sham groups before and after stimulation. This mean-level follow-up was guided by trial-level mixed-effects and localization results showing that stimulation-related modulation was strongest for negative prediction errors (Table S5). FRN amplitudes became more negative after stimulation in the active group, whereas no corresponding change was observed in the sham group.
